# DNA sequence differences are determinants of meiotic recombination outcome

**DOI:** 10.1101/568543

**Authors:** Simon D. Brown, Samantha J. Mpaulo, Mimi N. Asogwa, Marie Jézéquel, Matthew C. Whitby, Alexander Lorenz

## Abstract

Meiotic recombination is essential for producing healthy gametes, and also generates genetic diversity. DNA double-strand break (DSB) formation is the initiating step of meiotic recombination, producing, among other outcomes, crossovers between homologous chromosomes (homologs), which provide physical links to guide accurate chromosome segregation. The parameters influencing DSB position and repair are thus crucial determinants of reproductive success and genetic diversity. Using *Schizosaccharomyces pombe*, we show that the distance between sequence polymorphisms across homologs has a strong impact on meiotic recombination rate. The closer the sequence polymorphisms are to each other across the homologs the fewer recombination events were observed. In the immediate vicinity of DSBs, sequence polymorphisms affect the frequency of intragenic recombination events (gene conversions). Additionally, and unexpectedly, the crossover rate of flanking markers tens of kilobases away from the sequence polymorphisms was affected by their relative position to each other amongst the progeny having undergone intragenic recombination. A major regulator of this distance-dependent effect is the MutSα-MutLα complex consisting of Msh2, Msh6, Mlh1, and Pms1. Additionally, the DNA helicases Rqh1 and Fml1 shape recombination frequency, although the effects seen here are largely independent of the relative position of the sequence polymorphisms.

**Preamble:** Due to a mistake during analysis of a batch of Sanger sequencing reactions for Supplementary Figure S1, we erroneously stated that we found evidence for intragenic crossovers. We now show that intragenic crossovers are less likely than we initially thought. We sincerely apologize for our mishap and any inconvenience it might have caused. However, this does not affect the main conclusions of our paper, just how some of our results are interpreted. This new manuscript version has been amended accordingly.

## Introduction

Correct chromosome segregation during meiosis depends on pairing and physical connection of homologous chromosomes (homologs). Physical connections are established by the repair of programmed DNA double-strand breaks (DSBs) using the homolog rather than the sister chromatid as a template (i.e. interhomolog recombination) and by ensuring that interhomolog recombination intermediates are processed into crossovers (COs). The formation of DSBs by the transesterase Spo11 is thus a key step in initiating recombination during meiosis^1^. Regions of high-frequency Spo11 recruitment, and thus DSB formation, are called hotspots^2^. One of the best characterized category of hotspots are cAMP-responsive elements in *Schizosaccharomyces pombe*, created by point mutations in the *ade6* gene that represent binding sites for the Atf1-Pcr1 transcription factor^2,3^. These include the *ade6-M26* hotspot and its derivatives, which are defined by the DNA sequence heptamer 5’-ATGACGT-3’^3^. Although binding of Atf1-Pcr1 and the associated transcription already creates open chromatin at *M26*-like hotspots^3,4^, a very high frequency of meiotic recombination requires a conducive chromatin environment in a wider genomic context^5,6^.This network of parameters determines the overall level of DSB formation at a given genomic locus.

Following break formation, DSB ends are resected to initiate homologous recombination, which during meiosis follows either a Holliday junction/D-loop resolution or a synthesis-dependent strand annealing (SDSA) pathway^1,7^. As a repair template, either the sister chromatid or the homolog can be used^8^. Based on this, it has been suggested that the governance of meiotic recombination could be viewed as a two-tiered decision system^9^. The first decision being template choice (interhomolog vs. intersister recombination), and the second being how the recombination intermediate is resolved - i.e. the CO/non-crossover (NCO) decision. The template choice decision is mainly driven by meiosis-specific factors of the chromosome axis and by the meiotic recombinase Dmc1 supported by its mediators^8^. In budding yeast there is a basic understanding of how the interhomolog bias is established, although some mechanistic details still remain to be elucidated^10^. Since homologs are not necessarily identical on a DNA sequence level, a DSB end invading the homolog for repair can generate mismatch-containing heteroduplex DNA. Mismatches can be corrected by the mismatch repair system, consisting of the highly conserved MutS and MutL proteins^11^. Additionally, the MutS-MutL complex can also block strand invasion to avoid recombination between non-homologous sequences^11^. The CO/NCO-decision happens as the next step; here the decision is taken whether an already established interhomolog recombination intermediate is processed into a CO or a NCO. Determinants of the CO/NCO-decision are less well studied, but the DNA helicase/translocase FANCM (Fml1 in *Sz. pombe*) has been shown to limit CO formation in fission yeast and *Arabidopsis*^12,13^. RecQ-type DNA helicases perform a wide range of regulatory roles in homologous recombination, and one of which probably is the promotion of NCO formation during meiosis in various organisms^14–18^.

Here, we employ a series of meiotic recombination assays featuring intragenic markers at differently sized intragenic intervals and flanking intergenic markers to identify and characterize intrinsic determinants of template choice and CO/NCO-decision in fission yeast. We show that the relative positions of DNA sequence polymorphisms between homologs have a strong impact on recombination outcome, not only locally in the form of intragenic recombination (gene conversion), but also on the CO frequency between an up- and a downstream marker. The anti-recombinogenic activity of MutSα-MutLα factors, and of the DNA helicases Fml1 and Rqh1 modulate recombination outcome differentially when comparing various intragenic intervals.

## Results and Discussion

### Rationale of the meiotic recombination assay

Our meiotic recombination assay features intragenic markers (point mutations in the *ade6* gene) and flanking intergenic markers (*ura4^+^-aim2* and *his3^+^-aim*) (Fig. 1). This assay allows us to monitor various recombination outcomes: (I) intragenic recombination (gene conversion) events producing Ade^+^ recombinants, (II) crossovers (COs) between the flanking intergenic markers (*ura4^+^-aim2* and *his3^+^-aim*), and (III) the ratio of COs vs. non-crossovers (NCOs) among intragenic *ade6*^+^ recombination events (Fig. 1A). Changes in gene conversion and overall CO frequencies observed in this assay can be explained by an altered frequency of DSB formation at a given *ade6* mutant allele, or a change in repair template usage. The percentages of COs and NCOs among intragenic *ade6*^+^ recombination events are the genetic readout for the CO/NCO-decision, representing recombination intermediate processing after successful strand exchange between homologs. The intragenic events are most likely the result of gene conversions associated with COs or NCOs (non-reciprocal exchange of hereditary information).

**Figure 1.**
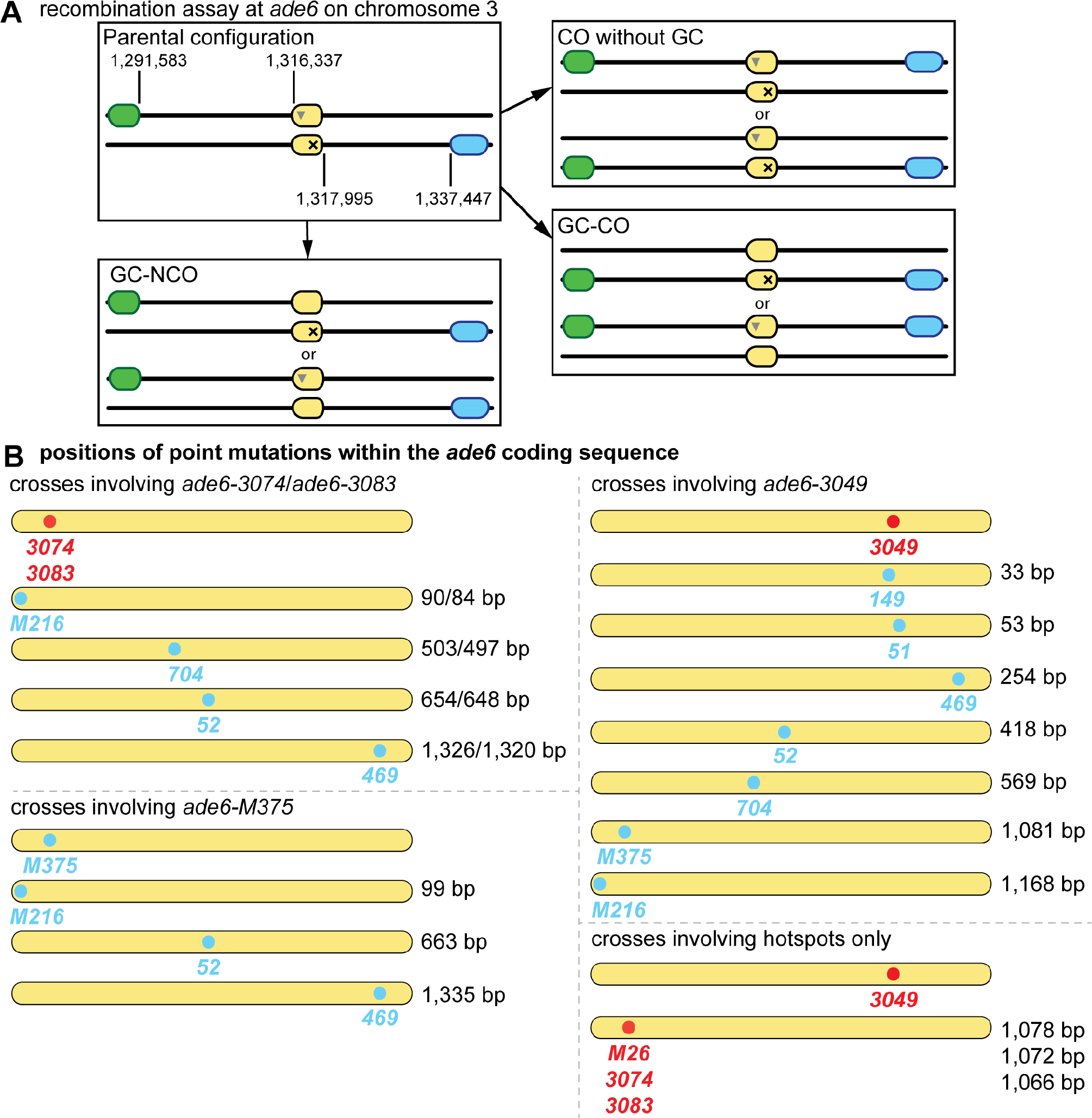
Meiotic recombination assay composed of *ade6* heteroalleles flanked by artificially introduced markers *ura4*^+^-*aim2* & *his3*^+^-*aim*. **(A)** Schematic showing the meiotic recombination assay at *ade6* (yellow) and its common outcomes. Ade^+^ recombinants can arise via gene conversion (GC) associated with a crossover (GC-CO) or a non-crossover (GC-NCO). The positions of *ade6*, and the artificially introduced markers *ura4*^+^-*aim2* (green) and *his3*^+^-*aim* (light blue) on chromosome 3 are indicated [in bps]. Positions of point mutations are shown as ▼ and ×. **(B)** Schematic of the *ade6* coding sequence indicating the point mutations and their positions (approximately to scale) used in the recombination assays, hotspots are indicated in red, and non-hotspots in light blue. The distance between the sequence polymorphisms across the homologs is indicated in relation to the given hotspot of each cross [in bp].

### The physical distance between point mutations of heteroalleles defines the frequency of intragenic recombination events and their associated CO/NCO ratio

Apart from absolute DSB levels, intragenic recombination frequency is also influenced by the distance between point mutations in a given chromosomal region^5,29–31^. Intragenic recombination in our assays (Fig. 1A) has so far been monitored using point mutations within the *ade6* coding sequence, which are at least 1kb apart^12,21,32^. We wondered whether the level of COs among intragenic recombination events also changes, when the distance between point mutations was decreased. Therefore, we selected a series of point mutations, which cover almost the complete length of the *ade6* coding sequence (Fig. 1B, Supplementary Table S1). These point mutants include the strong meiotic recombination hotspots *ade6-M26*, −*3074*, −*3083*, at the 5’ end of the gene and −*3049* at the 3’ prime end of the gene, and the non-hotspot alleles *ade6-M216*, −*M375*, −*704*, −*52*, −*149*, −*51*, and −*469* (Fig. 1B, Supplementary Table S1). All hotspots used here mimic a cAMP-response element, which creates a binding site for the Atf1-Pcr1 transcription factor; this in turn generates open chromatin^3,5^. It can be safely assumed that a given hotspot will receive the same amount of breakage independent of the *ade6* allele present on the homolog. This means that the differences seen in the combinations of one specific hotspot with various *ade6* alleles will depend on processes downstream of DSB formation. Indeed, the frequency of intragenic recombination positively correlates with the distance between the *ade6* alleles, when the same hotspot is used (Fig. 2A, black and grey lines). Recombination at the *ade6-M375* allele, which is at a similar position as the strong hotspot alleles *ade6-3074* & *ade6-3083*, is induced at an overall much lower level (Fig. 2A, green line), but appears to be the acceptor of genetic information when crossed to *ade6-469* (Fig. 2E), indicating that *ade6-M375* is somewhat more recombinogenic than *ade6-469*. Intragenic recombination frequency at *ade6-M375* shows a similar correlation with respect to distance between the DNA polymorphisms as crosses involving hotspots (Fig. 2A). Intragenic intervals of similar size containing the meiotic recombination hotspot alleles, *ade6-3083*, *ade6-3074*, or *ade6-3049*, and a non-hotspot allele produce equivalent intragenic recombination levels (Fig. 2A). Therefore, these hotspot alleles behave similarly in determining intragenic recombination frequency.

**Figure 2.**
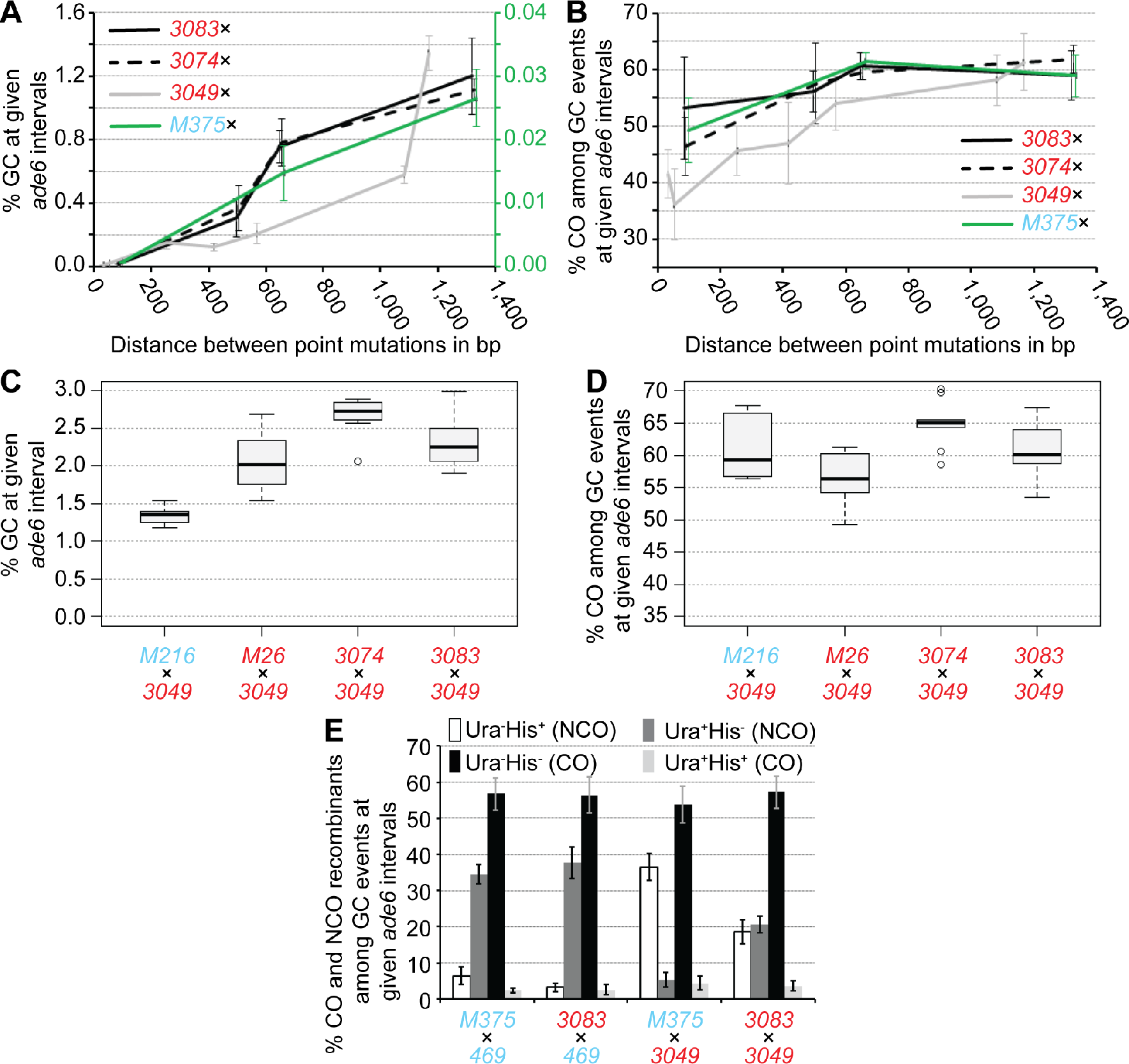
Physical distance between heteroalleles in *ade6* influences frequency of gene conversion (GC) and associated crossovers (COs). **(A)** Frequency of GC and **(B)** frequency of CO among GC events at *ade6* in wild type over distance between point mutations: crosses involving hotspot *ade6-3083* as black solid line, UoA110×UoA100 (*ade6-3083*×*ade6-M216*) (n=12), ALP733×UoA115 (*ade6-3083*×*ade6-704*) (n=12), ALP733×UoA119 (*ade6-3083*×*ade6-52*) (n=5), ALP733×ALP731 (*ade6-3083*×*ade6-469*) (n=20); crosses involving hotspot *ade6-3074* as black dashed line, UoA106×UoA100 (*ade6-3074*×*ade6-M216*) (n=12), UoA104×UoA115 (*ade6-3074*×*ade6-704*) (n=12), UoA104×UoA119 (*ade6-3074*×*ade6-52*) (n=6), UoA104×ALP731 (*ade6-3074*×*ade6-469*) (n=10); crosses involving hotspot *ade6-3049* as grey line, UoA122×UoA497 (*ade6-3049*×*ade6-149*) (n=6), UoA120×UoA463 (*ade6-3049*×*ade6-51*) (n=6), UoA120×ALP731 (*ade6-3049*×*ade6-469*) (n=31), UoA116×UoA123 (*ade6-3049*×*ade6-52*) (n=12), UoA112×UoA123 (*ade6-3049*×*ade6-704*) (n=12), ALP1541×UoA123 (*ade6-3049*×*ade6-M375*) (n=12), UoA99×UoA123 (*ade6-3049*×*ade6-M216*) (n=12); and crosses involving non-hotspot *ade6-M375* as green line – needs to be read from the green secondary y-axis in (A), UoA861×UoA100 (*ade6-M375*×*ade6-M216*) (n=6), ALP1541×UoA119 (*ade6-M375*×*ade6-52*) (n=6), ALP1541×ALP731 (*ade6-M375*×*ade6-469*) (n=16). **(C)** Frequency of GC and **(D)** frequency of CO among GC events at *ade6* in wild type crosses involving hotspot alleles only: FO1285×UoA123 (*ade6-M26*×*ade6-3049*) (n=12), UoA104×UoA123 (*ade6-3074*×*ade6-3049*) (n=9), and ALP733×UoA123 (*ade6-3083*×*ade6-3049*) (n=9); the hotspot×non-hotspot cross UoA99×UoA123 (*ade6-3049*×*ade6-M216*) (n=12) is shown for comparison. **(E)** Distribution of non-crossover (NCO; Ura^+^ His^−^ & Ura^−^ His^+^) and crossover (CO; Ura^+^ His^+^ & Ura^−^ His^−^) classes among Ade^+^ GC events in wild type (percentages in each class are shown as means ± Std. Dev.); ALP1541×ALP731 (n=16), ALP733×ALP731 (n=20), ALP1541×UoA123 (n=12), ALP733×UoA123 (n=9). n indicates the number of independent crosses. For details of data see Supplementary Table S2.

Intriguingly, these observations are also largely true for CO frequency among intragenic recombination events: The shorter an intragenic distance between polymorphisms is, the more likely an intragenic recombination event is resolved as a NCO (Fig. 2B). For crosses involving the hotspot alleles *ade6-3083* or *ade6-3074* the effect apparently tails off at intragenic distances >600 bp (Fig. 2B). Combining hotspot alleles on both homologs within a cross results in increased overall intragenic recombination rate compared with a hotspot × non-hotspot cross covering a similar intragenic distance between point mutations (Fig. 2C), in line with previous reports^33^. However, there is no notable difference in COs among intragenic recombination events (Fig. 2D). This indicates that the frequency of CO among intragenic recombination events is primarily a function of the distance between the *ade6* heteroalleles on the homologs.

The distribution of different NCO/CO classes amongst intragenic recombination events follows a pattern consistent with intragenic NCOs more likely being associated with the hotter allele. This means that the allele more likely to receive a DSB is the recipient of genetic information in the overwhelming majority of cases, which might represent a *bona fide* gene conversion event, e.g. the vast majority of Ade^+^ NCO events in the *ade6-3083*×*ade6-469* cross are Ura^+^ His^−^, because the *ade6-3083* allele is linked to the *ura4*^+^-*aim2* marker (Fig. 2E). If comparable hotspots are combined in a cross the two intragenic NCO classes occur with roughly equal frequency (Fig. 2E, compare cross *ade6-3083*×*ade6-3049* to crosses *ade6-3083*×*ade6-469* & *ade6-M375*×*ade6-3049*).

The observed distribution patterns also suggest that, at these long intragenic intervals, a subset of CO events could stem from the processing of one joint molecule, presumably a single Holliday junction^34^ or its precursors, positioned between the two *ade6* point mutations; in contrast to a gene conversion event being resolved as a CO. This idea makes the following prediction: If CO events among Ade^+^ recombinants (mostly Ura^−^ His^−^ genotypes) are created by processing of a joint molecule situated between the two *ade6* point mutations, then reciprocal Ura^+^ Ade^−^ His^+^ recombinants carrying the mutations of both *ade6* heteroalleles must exist. To test this, we sequenced the *ade6* locus from 32 Ura^+^ Ade^−^ His^+^ colonies from an *ade6-3083*×*ade6-469* cross. Based on the frequency of 0.677% Ura^−^ Ade^+^ His^−^ events among the total viable progeny in such a cross representing 8.375% of recombinants among all Ura^−^ His^−^ colonies (240 Ura^−^ His^−^ colonies among 2,969 total viable progeny, 8.083%), we would expect that 2-3 of the 32 Ura^+^ Ade^−^ His^+^ carry both the *3083* and the *469* mutation within the *ade6* locus, if all these events were generated by CO processing of a recombination internediate between the two heteroalleles. We did not observe any instances in which the *ade6* locus of Ura^+^ Ade^−^ His^+^ progeny harbored both mutations (Supplementary Fig. S1). Intragenic COs, if arising at all, are thus potentially only a minor cause such progeny among gene conversion, which are already relatively rare events. Rather, it is simple gene conversions at single loci, which are primarily generated by mismatch repair or DNA synthesis during DSB repair^35^, that are responsible.

### MutSα and MutLα are strong negative modulators of recombination frequency specifically at short intragenic intervals

Potential candidates for genetic pathways modulating recombination frequency at intragenic intervals of different lengths are MutS-MutL complexes, which bind to heteroduplex DNA and repair mismatches^11^. *Sz. pombe* has a streamlined nuclear mismatch repair system consisting of MutSα (Msh2-Msh6), MutSβ (Msh2-Msh3), and a single MutL (MutLα, Mlh1-Pms1); there is also a mitochondrial MutS protein called Msh1^36^. Importantly, the meiotic pro-crossover factors MutSγ (Msh4-Msh5), the meiosis-specific MutLγ component Mlh3, and Mlh2 – a MutLβ-homolog and a modulator of meiotic gene conversion tract length – are all missing in fission yeast^37,38^. This suggests that *Sz. pombe* is a suitable model to study the role of MutSα/β-MutLα during meiosis without potential crosstalk from MutSγ-MutLγ pro-crossover factors^39^.

At small intragenic intervals the absence of MutSα and/or MutLα causes a substantial increase in intragenic recombination frequency (Fig. 3A, Supplementary Fig. S2). This relationship shows an inverse correlation, i.e. the shorter the intragenic interval the higher the increase. This ranges from a ~70-fold increase at the *ade6*-*149*×*ade6*-*3049* (33 bp) interval, via a ~35-fold one at *ade6*-*3049*×*ade6*-*51* (53 bp), to a ~10-fold augmentation at the *ade6*-*M216*×*ade6*-*3083* (84 bp) interval (Fig. 3A, Supplementary Fig. S2). The *mutS*α mutants (*msh2-30*, *msh6*Δ) and the *mutL*α mutants (*mlh1*Δ, *pms1-16*) displayed similar frequencies of intragenic recombination to each other, and the *msh2-30 mlh1*Δ double mutant is not discernible from either single mutant (Fig. 3A), indicating that MutSα and MutLα work in the same pathway. Deleting *mutS*β (*msh3*) is of no consequence at the *ade6*-*M216*×*ade6*-*3083* interval (Fig. 3A; *p*=0.613 against wild type, two-tailed Mann-Whitney U), likely because all the *ade6* mutations tested are substitution mutations, and MutSβ only recognizes insertion/deletion loop mismatches larger than 2 nucleotides^11^. At larger intragenic intervals, there seems to be little or no role for MutSα-MutLα in limiting recombination events. A moderate, but mostly non-significant, tendency of lower intragenic recombination frequency can be observed (Fig. 3B, Supplementary Fig. S2). Altogether, these data show that MutSα-MutLα has a strong anti-recombinogenic role at small intragenic intervals, but seemingly no substantial role in determining recombination outcome at large intragenic intervals.

**Figure 3.**
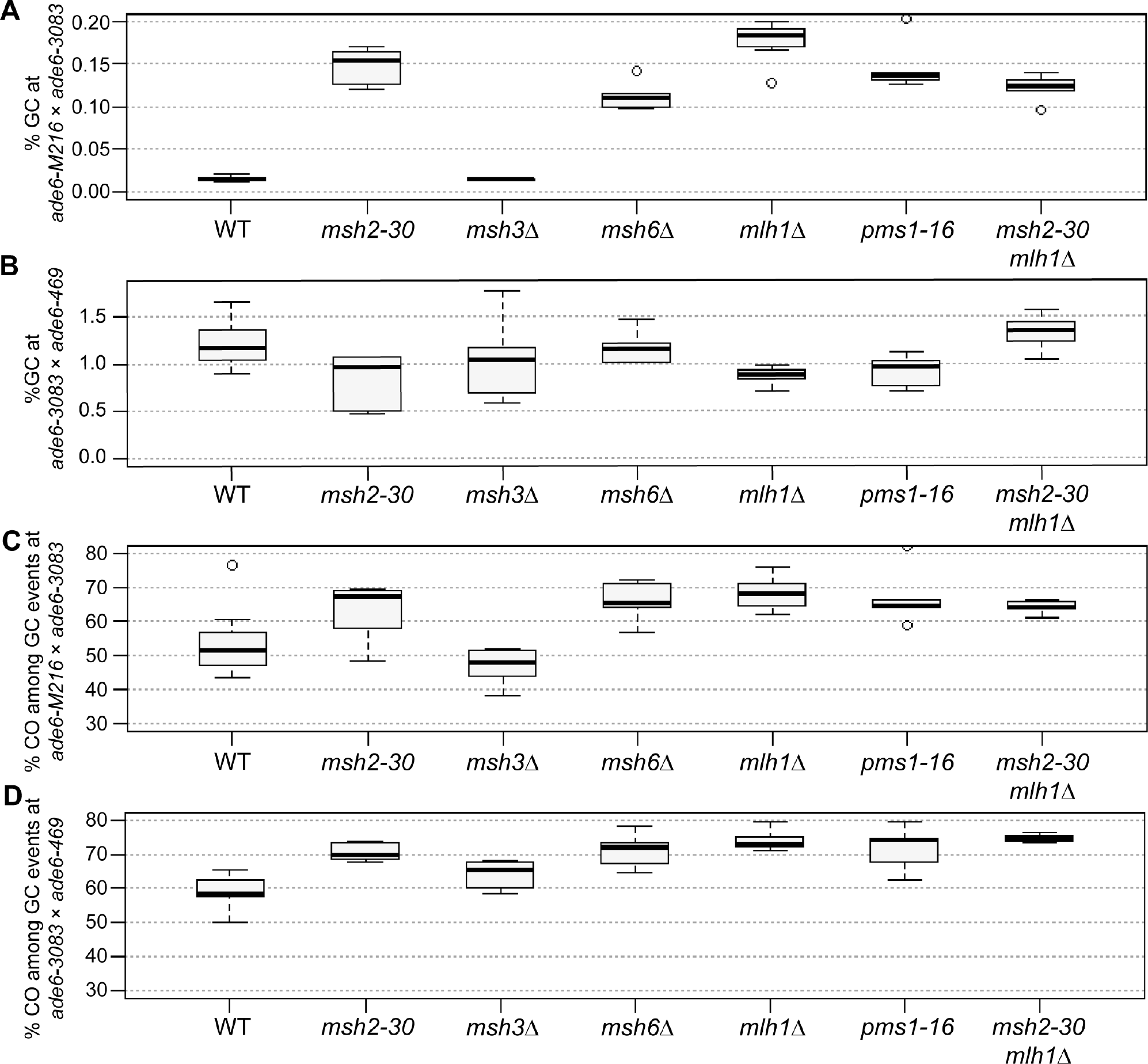
MutSα and MutLα, but not MutSβ, are major modulators of the gene conversion (GC) rate, and of the crossover (CO) frequency among GC events. **(A, B)** Frequency of GC in wild type (WT), *msh2*, *msh3*, *msh6*, *mlh1*, and *pms1* mutants **(A)** at the intragenic 84 bp interval *ade6-M216*×*ade6-3083*: UoA110×UoA100 (WT, n = 12), UoA478×UoA476 (*msh2-30*, n = 6), UoA494×UoA492 (*msh3*Δ, n = 6), UoA482×UoA480 (*msh6*Δ, n = 6), UoA364×UoA361 (*mlh1*Δ, n = 8), UoA407×UoA405 (*pms1-16*, n = 5), UoA828×UoA830 (*msh2-30 mlh1*Δ, n = 6); **(B)** at the intragenic 1,320 bp interval *ade6-3083*×*ade6-469*: ALP733×ALP731 (WT, n = 20), UoA477×UoA479 (*msh2-30*, n = 6), UoA493×UoA495 (*msh3*Δ, n = 6), UoA481×UoA483 (*msh6*Δ, n = 6), UoA362×UoA371 (*mlh1*Δ, n = 11), UoA406×UoA410 (*pms1-16*, n = 6), UoA827×UoA829 (*msh2-30 mlh1*Δ, n = 6). **(C, D)** Frequency of CO between *his3*^+^-*aim* and *ura4*^+^-*aim2* associated with GC events at *ade6* in wild type (WT), *msh2*, *msh3*, *msh6*, *mlh1*, and *pms1* mutants **(C)** at the intragenic 84 bp interval *ade6-M216*×*ade6-3083*: strains as in (A); **(D)** at the intragenic 1,320 bp interval *ade6-3083*×*ade6-469*: strains as in (B). n indicates the number of independent crosses. For details of data see Supplementary Table S3.

Mutating *mutS*α-*mutL*α genes increases CO frequency among gene conversion events (Fig. 3C-D, Supplementary Fig. S3) and/or changes the distribution of recombinant classes (Supplementary Fig. S4). Both long and short intragenic intervals involving the *ade6-3083* allele showed increases in associated CO frequency in comparison to wild type, albeit this trend was not statistically significant in all cases (Fig. 3C-D, Supplementary Fig. S3). This trend makes the share of COs among gene conversion events independent of the length of the interval (compare Fig. 2B with Fig. 3C-D, Supplementary Fig. S3).

Interestingly, there is also a substantial shift in CO classes among gene conversion events from mostly Ura^−^ His^−^ to mainly Ura^+^ His^+^ in *mutS*α-*mutL*α mutants at the short intervals *ade6*-*M216*×*ade6*-*3083* and *ade6*-*149*×*ade6*-*3049*, but not at the short *ade6*-*51*×*ade6*-*3049* interval (Supplementary Fig. S4A-C). At long intervals (*ade6*-*3083*×*ade6*-*469*, *ade6*-*M375*×*ade6*-*469*, *ade6*-*M216*×*ade6*-*3049*) this shift is not observed (Supplementary Fig. S4D-F). The change in CO classes among gene conversion events at the short intervals (*ade6*-*M216*×*ade6*-*3083* and *ade6*-*149*×*ade6*-*3049)* is not a consequence of selective survival or the formation of diploid or disomic spores, because *mutS*α-*mutL*α mutants have a spore viability similar to wild type, and the extent of the phenotype is the same in several different mutants (Supplementary Table S3). As with intragenic recombination frequency, the *mutS*β-deletion *msh3*Δ behaves just like wild type for CO outcome (Fig. 3C-D; *p*=0.439 against wild type, two-tailed Mann-Whitney U; Supplementary Fig. S4A and S4D).

The observed effects of different parental and recombinant classes amongst progeny having undergone a gene conversion event can be explained by envisioning a DSB 5’ or 3’ of a point mutation leading to a recombination intermediate (D-loop, Holliday junction), which will then be processed immediately at the break site, or ends up somewhat removed from the initial break site by multiple consecutive invasion steps, by branch migration, or both^40–42^. The genetic makeup of the progeny is, therefore, a compound result of processing distinct recombination intermediates in different ways. The genetic composition of wild-type and mutant progeny resulting from the meiotic recombination assays can be explained as different combinations of scenarios suggested previously^28^. For example, recombination between *ade6-3083* and *ade6-M216*, which gives rise to mainly Ura^−^ Ade^+^ His^+^ NCOs and Ura^−^ Ade^+^ His^−^ COs, may be explained by the model in Fig. 4A. In this model, a bias in favour of Ura^−^ Ade^+^ His^−^ COs stems from strand exchange/branch migration being constrained to within the region defined by the *ade6-3083* – *ade6-M216* interval and resolution of the recombination intermediate occurring by D-loop cleavage (Fig. 4A, C). Ura^-^ Ade^+^ His^+^ NCOs and additional Ura^−^ Ade^+^ His^−^ COs come from HJ resolution (Fig. 4A, C).

**Figure 4.**
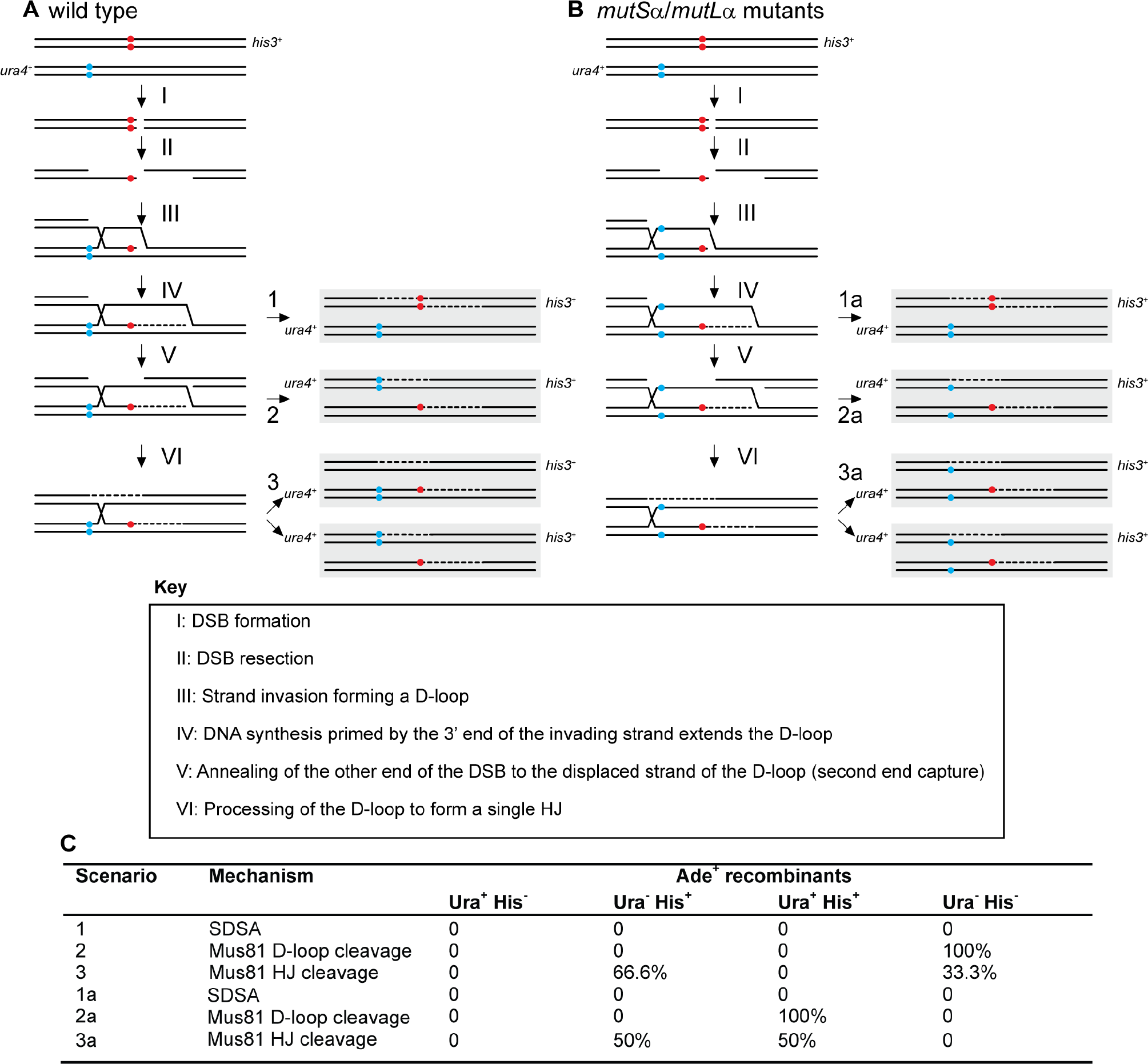
Possible scenarios for CO/NCO recombination events creating Ade^+^ progeny from crosses with different *ade6* heteroalleles and *ura4^+^-aim2* and *his3^+^-aim* as flanking markers. **(A, B)** The two black lines represent double-stranded DNA of one chromatid; chromatids not involved in the depicted recombination event are omitted for clarity. Positions of the hotspot and non-hotspot alleles are indicated in red and light blue, respectively. **(A)** Predominant situation in wild type, where Ade^+^ CO recombinants are mostly Ura^−^ His^−^. **(B)** Situation explaining the Ura^+^ Ade^+^ His^+^ progeny observed in some *mutS*α-*mutL*α mutant crosses. Extensive branch migration and/or multiple invasion events could cause the D-loop or Holliday Junction (HJ) eventually being established left of the non-hotspot allele. Subsequent processing will generate Ura^+^ Ade^+^ His^+^ CO progeny at a high frequency. **(C)** Frequency of possible recombination outcomes in crosses involving two *ade6* heteroalleles and flanking markers (*ura4*^+^*-aim2* and *his3*^+^*-aim*) as shown in (A) and (B).

We also considered whether this alteration of recombination outcome at *ade6-M216*×*ade6-3083* in *mutS*α-*mutL*α mutants, which leads to relatively few Ade^+^ His^−^ Ura^−^ COs and a big increase in the proportion of Ade^+^ His^+^ Ura^+^ COs (Fig. 3, Supplementary Fig. S4A), might have something to do with the complexity of the *ade6-3083* allele. This allele consists of multiple substitution mutations and can potentially form a C/C-mismatch in the heteroduplex DNA during strand exchange that is less efficiently repaired during meiosis than other mismatches^43^. However, as mentioned above, a moderate shift of CO recombinant classes among intragenic events can also be seen at another small interval, *ade6*-*149*×*ade6*-*3049* (Supplementary Fig. S4B). Unlike *ade6-3083*, *ade6*-*3049* contains only a single nucleotide difference (Supplementary Table S1) and, therefore, the complexity of a given *ade6* allele is unlikely to be the critical factor affecting the shift in CO recombinant class. This is complicated by the fact that a third small interval, *ade6*-*51*×*ade6*-*3049*, does not show this shift between CO recombinant categories, similar to long intervals (Supplementary Fig. S4C-F). We think that a deficit in heteroduplex rejection and mismatch repair, caused by loss of *msh2*, could results in strand exchange/branch migration extending beyond the non-hotspot mutation (i.e. *ade6-M216* or *ade6-149*) prior to D-loop cleavage/HJ resolution, with the base-pair mismatches in the recombinant chromosomes remaining unrepaired. Together, these altered features could explain the increase in Ade^+^ His^+^ Ura^+^ COs at the *ade6-M216*×*ade6-3083* and *ade6*-*149*×*ade6*-*3049* intervals in *mutS*α-*mutL*α mutant crosses (Fig. 4B, C). However, why *ade6*-*51*×*ade6*-*3049* would not show this behavior remains unclear; potentially the positioning of the DSBs in relation to the hotspot mutations could play a role here.

Recombination outcome in a *msh2*Δ in *S. cerevisiae* has also been shown to be more complex than in wild type^44,45^. Intriguingly, in *S. cerevisiae* the action of Msh2 seems to be restricted to class I COs, which are subjected to CO interference, whereas Mus81-dependent class II COs are unchanged in *msh2*Δ^45^. *Sz. pombe* operates only a class II CO pathway via Mus81-processing, completely lacking a class I CO pathway. Nevertheless, the absence of Msh2 in fission yeast has a profound effect on CO frequency, and the way recombination intermediates are processed (Fig. 3, Supplementary Fig. S4).

### Fml1 is a negative modulator of CO frequency among gene conversion events independent of the distance between point mutations

The DNA helicases, Fml1 and Rqh1, are also prime candidates for modulating recombination frequency at intragenic intervals of different lengths^12,46^. However, Fml1 apparently does not modulate gene conversion levels, as at all intragenic intervals tested, *fml1*Δ is similar to wild type (Fig. 5A-B, Supplementary Fig. S5A). In contrast, the RecQ-family DNA helicase Rqh1 is required for wild-type levels of gene conversion^12^. The deletion of *rqh1* reduces gene conversion frequency to about a third of wild-type percentage at short (*ade6*-*M216*×*ade6*-*3083*, *ade6*-*3049*×*ade6*-*469*) intervals, and to about a tenth of wild-type frequency at the long *ade6*-*3083*×*ade6*-*469* interval (Fig. 5A-B, Supplementary Fig. S5).

**Figure 5.**
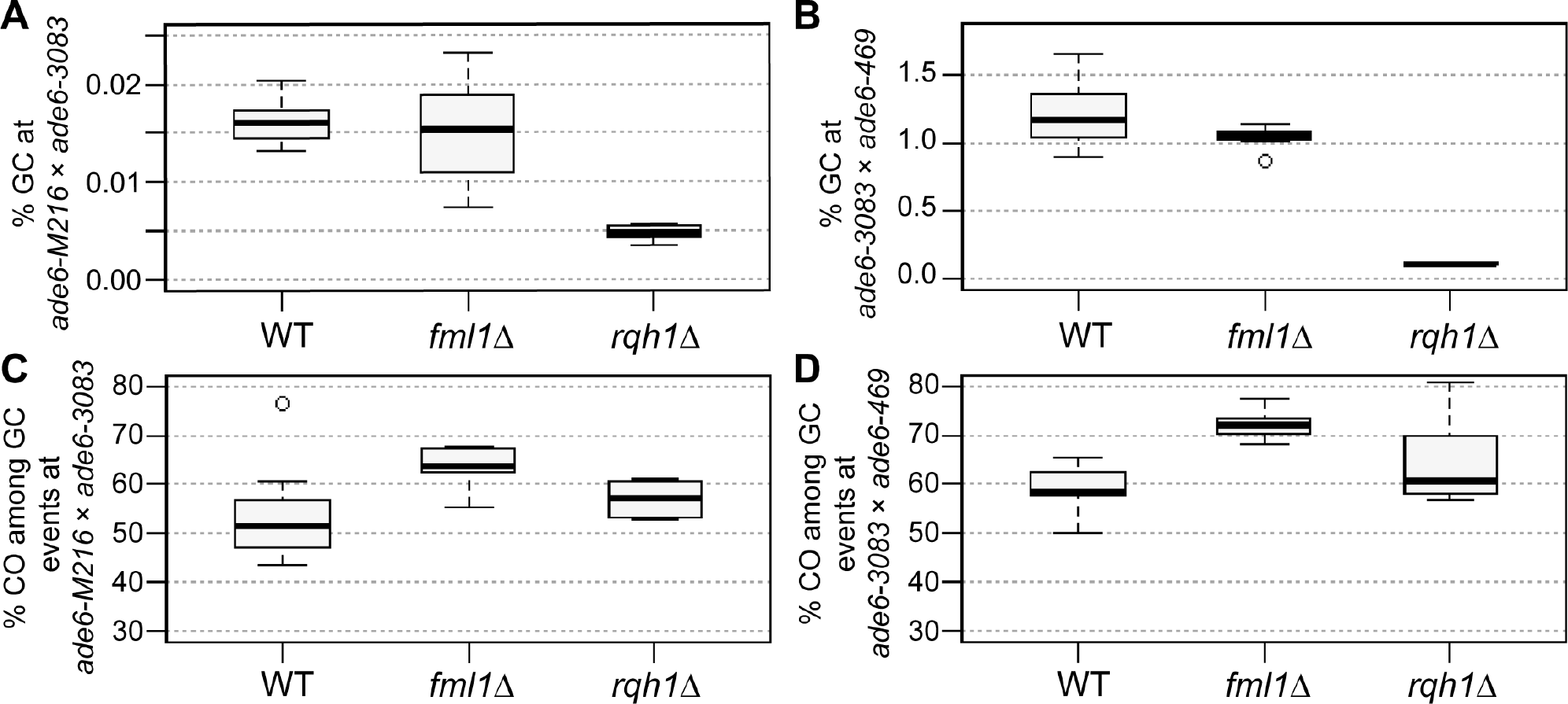
The RecQ-family helicase Rqh1, but not the FANCM-type helicase Fml1, is a major modulator of the gene conversion (GC) rate. Rqh1 and Fml1 are major modulators of crossover (CO) frequency among GC events. Frequency of GC in WT, *fml1*, and *rqh1* deletions **(A)** at the intragenic 84 bp interval *ade6-M216*×*ade6-3083*: UoA110×UoA100 (WT, n = 12), UoA450×UoA447 (*fml1*Δ, n = 9), UoA502×UoA499 (*rqh1*Δ, n = 6); **(B)** at the intragenic 1,320 bp interval *ade6-3083*×*ade6-469*: ALP733×ALP731 (WT, n = 20), ALP1133×MCW4718 (*fml1*Δ, n = 15), ALP781×ALP780 (*rqh1*Δ, n = 10). Frequency of CO between *his3*^+^-*aim* and *ura4*^+^-*aim2* associated with GC events at *ade6* in WT, *fml1*, and *rqh1* deletions **(C)** at the intragenic 84 bp interval *ade6-M216*×*ade6-3083*: strains as in (A); **(D)** at the intragenic 1,320 bp interval *ade6-3083*×*ade6-469*: strains as in (B). n indicates the number of independent crosses. For details of data see Supplementary Table S3.

As with long intervals^12^, *fml1*Δ results in a ~10 percentage point increase of CO frequency among gene conversion events at short intervals (Fig. 5C-D, Supplementary Fig. S5). The absence of Rqh1 induces moderate increases in CO levels among gene conversion events at the 84 bp *ade6*-*M216*×*ade6*-*3083* and the 1,320 bp *ade6*-*3083*×*ade6*-*469* interval, which are not statistically significant (Fig. 5C-D). However, at the 254 bp *ade6*-*3049*×*ade6*-*469* interval CO frequency among *ade6*^+^ events is raised by 17 percentage points in *rqh1*Δ (*p*=3.72×10^−9^ against wild type, two-tailed Mann-Whitney U) (Supplementary Fig. S5). The *ade6-3083* allele contains multiple point mutations and thus represents a more complex situation than the *ade6-3049* allele, which only harbors a single point mutation. Fml1 can seemingly drive NCO pathway(s) independently of the complexity of the underlying DNA sequence, because it has the same effect in crosses with complex and single-mutation alleles. In contrast, Rqh1 can apparently fulfill this role only at the simple *ade6-3049* allele.

In *Sz. pombe* Fml1 has been shown to specifically limit CO formation during the late CO/NCO-decision^12^. Fml1 acts as a promotor of NCOs, likely by driving late recombination intermediates into the SDSA pathway, after strand invasion and DNA synthesis has happened. In accordance with this, absence of *fml1* leads to an increase in CO among Ade^+^ gene conversion events, but has little effect on intragenic recombination itself (Fig. 5, Supplementary Fig. S5)^12^. This role is independent of the size of the intragenic interval, with Fml1 driving 10-12% of NCO recombination in any case.

The deletion of *rqh1* has a very strong meiotic phenotype, leading to reductions in intragenic recombination, CO, and spore viability (Fig. 5, Supplementary Fig. S5). This on its own would indicate an early role in promoting strand exchange and/or DSB resection, but then Rqh1 additionally is capable of promoting NCO formation among *ade6*^+^ events at some intragenic intervals during later stages of recombination (Fig.5, Supplementary Fig. S5). Most likely this is due to Rqh1 actually performing the following functions: (I) promotion of interhomolog recombination events, probably in cooperation with Rad55-57 and Rlp1-Rdl1-Sws1, but independently of Sfr1-Swi5,^28^ potentially also by providing longer resection tracts^47^; (II) dismantling D-loops, this enables the release of break ends to search for homology elsewhere, starts cycles of multiple consecutive invasion steps, and provides opportunities for Fml1 to drive NCO formation via SDSA; and (III) branch migration of established D-loops and Holliday junctions, thereby promoting heteroduplex DNA formation further away from the break site^46^.

Overall, these data show that Fml1 has likely no role in modulating gene conversion levels, but drives NCO formation downstream after successful strand invasion and DNA synthesis. Rqh1 promotes intragenic recombination, but also has moderate anti-recombinogenic activity in CO formation among gene conversion events.

In conclusion, factors directly involved in generating CO and NCO recombinants during meiosis have been identified and characterized in recent years^12–15,21^, and several inroads have been made in understanding how template choice is regulated and executed during meiotic recombination^10,28^. However, we still only have a basic understanding of how underlying DNA sequence polymorphisms influence meiotic recombination outcomes. This is critically important for understanding recombination event distribution in natural populations, where any two parental genomes will be littered with sequence polymorphisms. Here, we demonstrate that specific DNA sequence differences between the two homologs strongly impact on which outcome is achieved, and that this is largely driven by the action of the MutS-MutL complex. This highlights the importance of the interplay between *cis*- and *trans*-factors in shaping the genetic diversity of a given population.

## Material and methods

### Bacterial and yeast strains and culture conditions

*E. coli* strains were grown on LB and SOC media – where appropriate containing 100 µg/ml Ampicillin^19^. Competent cells of *E. coli* strains NEB10^®^-beta (New England BioLabs Inc., Ipswich, MA, USA), and XL1-blue (Agilent Technologies, Santa Clara, CA, USA) were transformed following the protocols provided by the manufacturers. *Sz. pombe* strains used for this study are listed in Supplementary Table S4. Yeast cells were cultured on yeast extract (YE), and on yeast nitrogen base glutamate (YNG) agar plates containing the required supplements (concentration 250 mg/l on YE, 75 mg/l on YNG). Crosses were performed on malt extract (ME) agar containing supplements at a final concentration of 50 mg/l^20^.

Different *ade6* hotspot and non-hotspot sequences (Supplementary Table S1) were introduced by crossing the respective mutant *ade6* strain with *ade6*^+^ strains carrying the *ura4*^+^ and *his3*^+^ <u>a</u>rtificially <u>i</u>ntroduced <u>m</u>arkers (aim) (UoA95, UoA96, UoA97, UoA98)^21^. The point mutations in the *ade6* alleles were verified by Sanger DNA sequencing (Source BioScience, Nottingham, UK) (Supplementary Table S1).

Using an established marker swap protocol^22^ the *natMX6*-marked *rqh1*Δ-*G1* was derived from an existing *rqh1*Δ∷*kanMX6* allele^23^, creation of the *natMX6*-marked *pms1-16* insertion mutant allele has been described previously^24^.

Marker cassettes to delete *msh3*, and *msh6*, and to partially delete *msh2* were constructed by cloning targeting sequences of these genes into pFA6a-*kanMX6*, pAG25 (*natMX4*), and pAG32 (*hphMX4*), respectively, up- and downstream of the dominant drug resistance marker^25,26^. The targeting cassettes were released from plasmids (pALo130, pALo132, pALo134) generated for this purpose by a restriction digest, and transformed into the strains FO652 (*msh2* and *msh6*) and ALP729 (*msh3*). For specifics of strain and plasmid construction, please refer to Supplementary Materials. Plasmid sequences are available on figshare (https://dx.doi.org/10.6084/m9.figshare.6949274).

Transformation of yeast strains was performed using an established lithium-acetate procedure^27^. All plasmid constructs were verified by DNA sequencing (Source BioScience plc, Nottingham, UK).

All DNA modifying enzymes (high-fidelity DNA polymerase Q5, restriction endonucleases, T4 DNA ligase) were supplied by New England BioLabs. Oligonucleotides were obtained from Sigma-Aldrich Co. (St. Louis, MO, USA).

### Genetic and molecular assays

Determination of spore viability by random spore analysis and the meiotic recombination assay have been previously described in detail^20,21^.

To test whether intragenic COs exist, genomic DNA of Ura^+^ Ade^−^ His^+^ progeny from an *ade6-3083*×*ade6-469* (ALP733×ALP731) cross was used to PCR-amplify the *ade6* locus (oligonucleotides oUA219 5’-AAAGTTGCATTTCACAATGC-3’ and oUA66 5’-GTCTATGGTCGCCTATGC-3’) for Sanger sequencing (Eurofins Scientific, Brussels, Belgium) with oUA219, oUA66, or nested oligonucleotides oUA779 5’-CTCATTAAGCTGAGCTGCC-3’ and oUA780 5’-AAGCTCTCCATAGCAGCC-3’.

### Data presentation and Statistics

Raw data is available on figshare (https://dx.doi.org/10.6084/m9.figshare.6949274). Line graphs were produced using Microsoft Excel 2016 (version 16.0.4638.1000, 32-bit). Box-and-whisker plots were created in R (version i386, 3.0.1) (http://www.r-project.org/) using the standard settings of the boxplot() function^28^. The lower and upper ‘hinges’ of the box represent the first and third quartile, and the bar within the box indicates the median (=second quartile). The ‘whiskers’ represent the minimum and maximum of the range, unless they differ more than 1.5-times the interquartile distance from the median. In the latter case, the borders of the 1.5-times interquartile distance around the median are indicated by the ‘whiskers’ and values outside this range (‘outliers’) are shown as open circles. R was also used to compute Kruskal-Wallis test and Tukey’s Honest Significant Differences employing the kruskal.test() and TukeyHSD() functions, respectively. Mann-Whitney U tests were performed as previously described^28^.

## Acknowledgments

We are grateful to Jürg Bähler, Edgar Hartsuiker, Franz Klein, Jürg Kohli, Josef Loidl, Kim Nasmyth, Fekret Osman, Gerald R. Smith, Walter W. Steiner, and the National BioResource Project (NBRP) Japan for providing materials, and to C. Bryer, A. Mehats, and H. Rickman for technical assistance. This work was supported by the Biotechnology and Biological Sciences Research Council UK (BBSRC) [grant numbers BB/F016964/1, BB/M010996/1], the University of Aberdeen (College of Life Sciences and Medicine Start-up grant to AL), and the Wellcome Trust (Programme grant to MCW) [grant number 090767/Z/09/Z].

## Author Contributions

AL and MCW conceived this study. AL, SDB, SJM, MNA, and MJ conducted the experiments. AL drafted the manuscript. All authors read, revised and approved the manuscript.

## Competing Interests

The authors declare no competing interests

## Notes

#### Summary of Updates

Due to a mistake during analysis of a batch of Sanger sequencing reactions for Supplementary Figure S1, we erroneously stated that we found evidence for intragenic crossovers. We now show that intragenic crossovers are less likely than we initially thought. We sincerely apologize for our mishap and any inconvenience it might have caused. However, this does not affect the main conclusions of our paper, just how some of our results are interpreted. This new manuscript version has been amended accordingly.

https://dx.doi.org/10.6084/m9.figshare.6949274

