## Supplementary Methods, Figures & Tables S1 & S4 for "DNA sequence differences are determinants of meiotic recombination outcome"

**Specifics of yeast strain and plasmid construction**

The open reading frame of *msh2* (*SPBC19G7.01c*) overlaps with the open reading frame of *cwf14*, to avoid potentially affecting Cwf14 expression and function only a small portion of the 5' end of *msh2* was deleted. A targeting cassette for *msh2* was constructed by cloning an upstream flanking sequence of *msh2* (PCR using oligonucleotides oUA47 5'-AATTAACAGCTGCTTTAGAAAGTTCCCACC-3' and oUA48 5'-AATTAAGGATCCGCATTCGAACTATTAAACACC-3' on genomic DNA of ALP1594) digested with *Pvu*II and *Bam*HI into pAG32[1] linearized with *Pvu*II and *Bam*HI. The resulting plasmid (pALo129) was linearized by digesting with *Sac*I and *Spe*I and a part of the coding sequence of *msh2* (PCR using oligonucleotides oUA49 5'-AATTAAGAGCTCGTTTTCTAGGAATTTTACGTTGC-3' and oUA50 5'-AATTAAACTAGTCAAGTTCAACATCTCGAGC-3' on genomic DNA of ALP1594) digested with *Sac*I and *Spe*I was inserted by standard cloning to give pALo130. The transformation cassette was released by a *Pvu*II-*Spe*I digest and transformed into the standard lab strain FO652. This construct removes the 242 bps at the very 5' end of the *msh2* coding sequence plus an additional 299 bps upstream of the Start codon. Correct integration was monitored by selection for hygromycin B resistance and verified by PCR; all strains carrying the *msh2-30*::*hphMX4* insertion mutation are derived by crossing from the original transformant (UoA459).

A deletion cassette for *msh3* (*SPAC8F11.03*) was constructed by cloning an upstream flanking sequence of *msh3* (PCR using oligonucleotides oUA51 5'-AATTAACAGCTGCACGATGTAAAGAGTAGC-3' and oUA52 5'-AATTAAGGATCCGCTCAACATAGATTTGTAACG-3' on genomic DNA of ALP1594) digested with *Pvu*II and *Bam*HI into pAG25[1] linearized with *Pvu*II and *Bam*HI. The resulting plasmid (pALo131) was linearized by digesting with *Sac*I and *Spe*I and a downstream flanking sequence of *msh3* (PCR using oligonucleotides oUA53 5'-AATTAAGAGCTCGAAGAAATCTGAGAGAGAGC-3' and oUA54 5'-AATTAAACTAGTCTAAAAGAGCAGAGCAAACC-3' on genomic DNA of ALP1594) digested with *Sac*I and *Spe*I was inserted by standard cloning to give pALo132. The transformation cassette was released by a *Pvu*II-*Spe*I digest and transformed into the standard lab strain ALP729. This construct almost completely removes the *msh3* coding sequence: at the 5' end an additional 37 bps upstream of the Start codon are deleted, and at the 3' end the last 12 bps of the coding sequence are retained. Correct integration was monitored by selection for CLONNAT-resistance and verified by PCR; all strains carrying the *msh3Δ-32*::*natMX4* deletion are derived by crossing from the original transformant (UoA460).

A deletion cassette for *msh6* (*SPCC285.16c*) was constructed by cloning an upstream flanking sequence of *msh6* (PCR using oligonucleotides oUA55 5'-AATTAACAGCTGTTCTCTTTGCTGGTTTC-3' and oUA56 5'-AATTAAGGATCCGAACAAGTGTTGGTTTTGG-3' on genomic DNA of ALP1594) digested with *Pvu*II and *Bam*HI into pFA6a-*kanMX6*[2] linearized with *Pvu*II and *Bam*HI. The resulting plasmid (pALo133) was linearized by digesting with *Sac*I and *Spe*I and a downstream flanking sequence of *msh6* (PCR using oligonucleotides oUA57 5'-AATTAAGAGCTCACCTCTCATACTTGATTCG-3' and oUA58 5'-AATTAAACTAGTCGTTACGAATAATGGAACG-3' on genomic DNA of ALP1594) digested with *Sac*I and *Spe*I was inserted by standard cloning to give pALo134. The transformation cassette was released by a *Pvu*II-*Spe*I digest and transformed into the standard lab strain FO652. This construct almost completely removes the *msh6* coding sequence: at the 5' end an additional 4 bps upstream of the Start codon are deleted, and at the 3' end the last 9 bps of the coding sequence are retained. Correct integration was monitored by selection for G418-resistance and verified by PCR; all strains carrying the *msh6Δ-34*::*kanMX6* deletion are derived by crossing from the original transformant (UoA461).

All plasmid constructs were verified by DNA sequencing (Source BioScience plc, Nottingham, UK). DNA modifying enzymes (high-fidelity DNA polymerase Q5, restriction endonucleases, T4 DNA ligase) were supplied by New England BioLabs. Oligonucleotides were obtained from Sigma-Aldrich Co. (St. Louis, MO, USA).

Plasmid sequences are available online as supporting material (https://dx.doi.org/10.6084/m9.figshare.6949274).

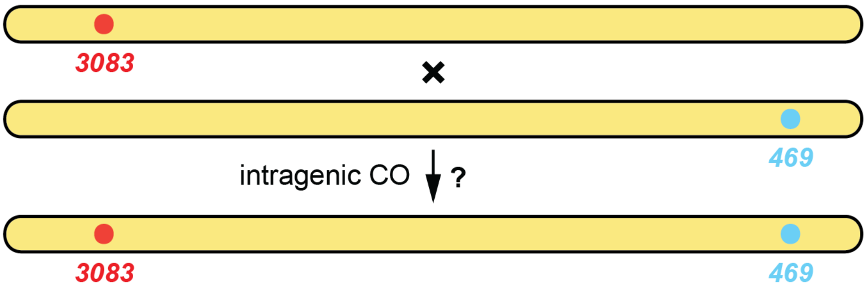


*ade6* sequence of 32 Ade⁻ Ura⁺ His⁺ progeny from cross ALP733 (*ade6-3083*) × ALP731 (*ade6-469*)

| colony number | 5' end | 3' end |
|---|---|---|
| 1 | ***3083*** | **wt** |
| 2 | **wt** | *469* |
| 3 | ***3083*** | **wt** |
| 4 | **wt** | ***469*** |
| 5 | **wt** | ***469*** |
| 6 | **wt** | ***469*** |
| 7 | **wt** | ***469*** |
| 8 | ***3083*** | **wt** |
| 9 | **wt** | *469* |
| 10 | ***3083*** | **wt** |
| 11 | **wt** | *469* |
| 12 | **wt** | ***469*** |
| 13 | ***3083*** | **wt** |
| 14 | **wt** | ***469*** |
| 15 | ***3083*** | **wt** |
| 16 | ***3083*** | **wt** |
| 17 | ***3083*** | **wt** |
| 18 | ***3083*** | **wt** |
| 19 | ***3083*** | **wt** |
| 20 | **wt** | ***469*** |
| 21 | **wt** | *469* |
| 22 | ***3083*** | **wt** |
| 23 | **wt** | *469* |
| 24 | ***3083*** | **wt** |
| 25 | **wt** | *469* |
| 26 | ***3083*** | **wt** |
| 27 | ***3083*** | **wt** |
| 28 | **wt** | *469* |
| 29 | ***3083*** | **wt** |
| 30 | ***3083*** | **wt** |
| 31 | **wt** | *469* |
| 32 | **wt** | *469* |

**Supplementary Figure S1.** Intragenic COs between the *3083* and the *469* point mutations in *ade6* could not be confirmed. The *ade6* locus was sequenced in 32 Ade- Ura+ His+ colonies from an *ade6-3083*×*ade6-469* (ALP733×ALP731) cross, no instances carrying both mutations were recorded. wt (wild type), *3083*, and *469* in bold indicate the status of the sequence confirmed by Sanger sequencing at the 5' and 3' ends, respectively. At the 3' end, the presence of *469* was assumed in some cases (not bold, black) based on the colony being Ade⁻ and having a wt sequence at the 5' end.

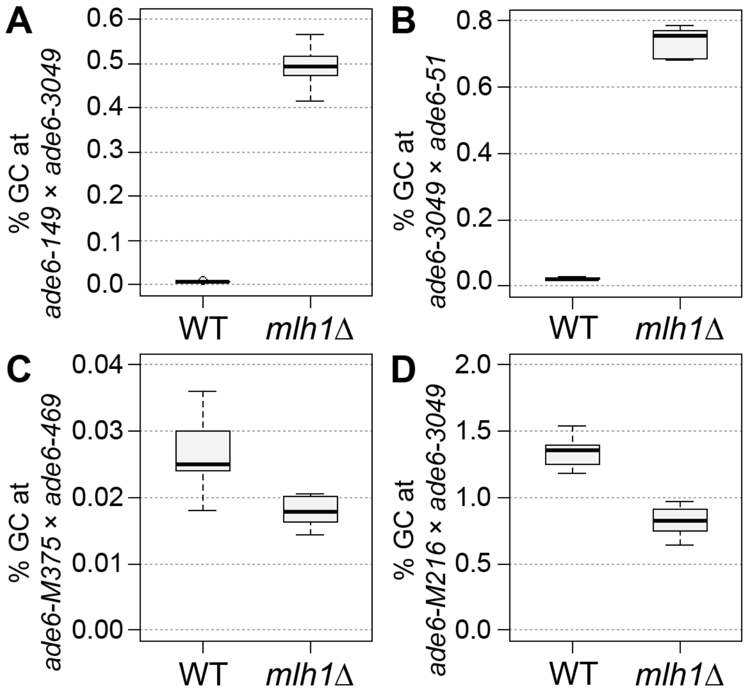


**Supplementary Figure S2.** MutLα is a major modulator of gene conversion (GC) rate. Frequency of GC in wild type (WT), and *mlh1*Δ. **(A)** at the intragenic 33 bp interval *ade6-149*×*ade6-3049*: UoA122×UoA497 (WT, n = 6), UoA368×UoA512 (*mlh1*Δ, n = 6); **(B)** at the intragenic 53 bp interval *ade6-3049*×*ade6-51*: UoA120×UoA463 (WT, n = 6), UoA366×UoA511 (*mlh1*Δ, n = 6); **(C)** at the intragenic 1,335 bp interval *ade6-M375*×*ade6-469*: ALP1541×ALP731 (WT, n = 16), UoA510×UoA371 (*mlh1*Δ, n = 6); **(D)** at the intragenic 1,168 bp interval *ade6-M216*×*ade6-3049*: UoA99×UoA123 (WT, n = 12), UoA368×UoA361 (*mlh1*Δ, n = 12); n indicates the number of independent crosses. For details of data see Supplementary Table S3.

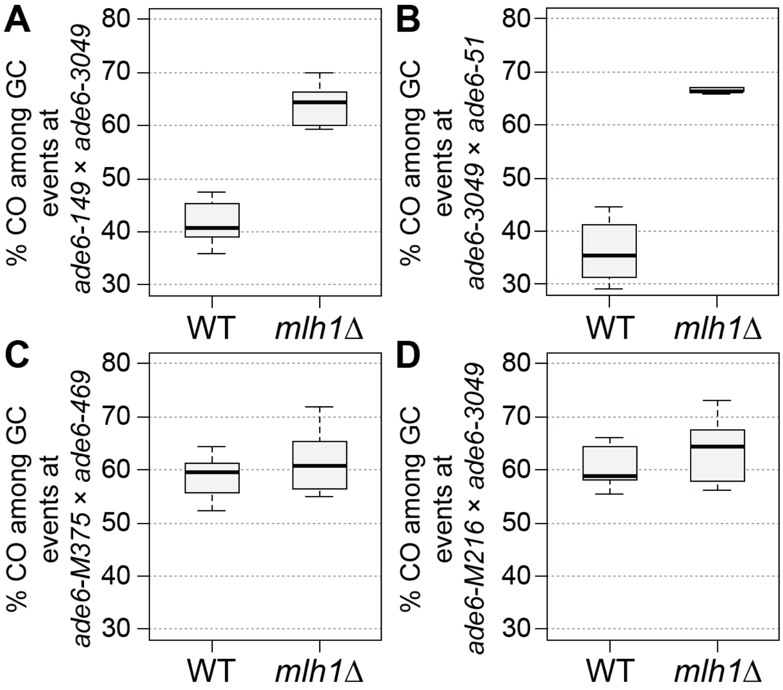


**Supplementary Figure S3.** MutLα is a major modulator of crossover (CO) frequency among gene coversion (GC) events. Frequency of CO between *his3*$^{+}$*-aim* and *ura4*$^{+}$*-aim2* associated with GC events at *ade6* in wild type (WT), and *mlh1*Δ. **(A)** at the intragenic 33 bp interval *ade6-149*×*ade6-3049*: UoA122×UoA497 (WT, n = 6), UoA368×UoA512 (*mlh1*Δ, n = 6); **(B)** at the intragenic 53 bp interval *ade6-3049*×*ade6-51*: UoA120×UoA463 (WT, n = 6), UoA366×UoA511 (*mlh1*Δ, n = 6); **(C)** at the intragenic 1,335 bp interval *ade6-M375*×*ade6-469*: ALP1541×ALP731 (WT, n = 16), UoA510×UoA371 (*mlh1*Δ, n = 6); **(D)** at the intragenic 1,168 bp interval *ade6-M216*×*ade6-3049*: UoA99×UoA123 (WT, n = 12), UoA368×UoA361 (*mlh1*Δ, n = 12); n indicates the number of independent crosses. For details of data see Supplementary Table S3.

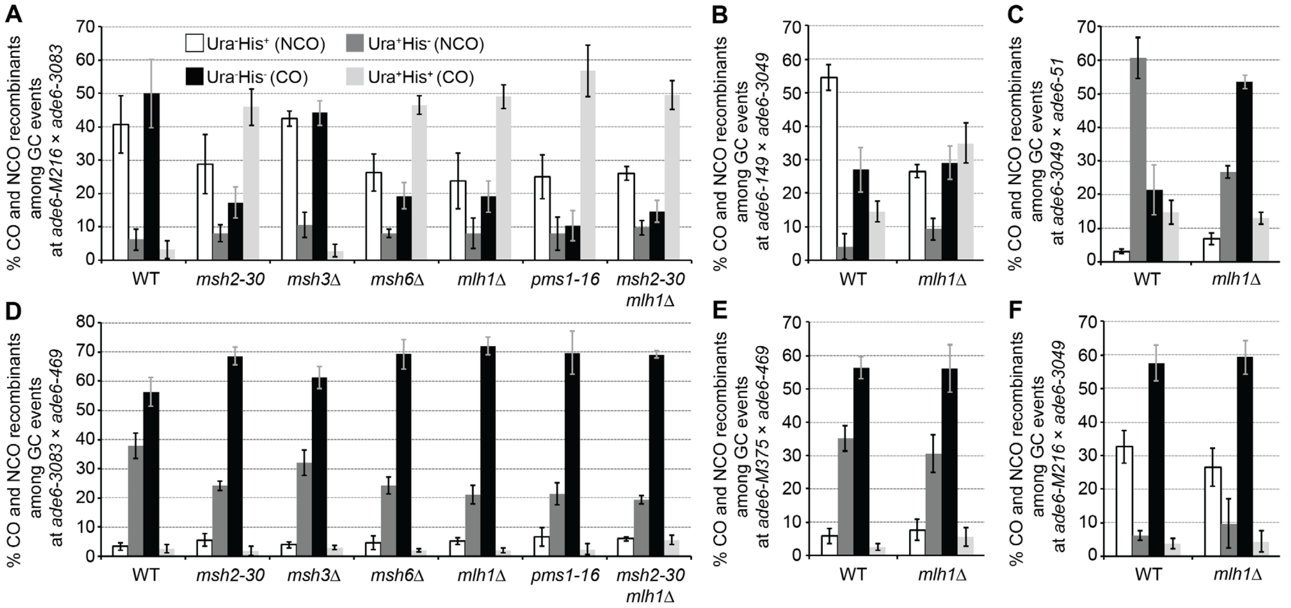


**Supplementary Figure S4.** Distribution of non-crossover (NCO; Ura$^{+}$ His$^{-}$ & Ura$^{-}$ His$^{+}$) and crossover (CO; Ura$^{+}$ His$^{+}$ & Ura$^{-}$ His$^{-}$) classes among Ade$^{+}$ gene conversion (GC) events in wild type (WT), *msh2*, *msh3*, *msh6*, *mlh1*, and *pms1* mutants (percentages in each class are shown as means ± Std. Dev.. **(A)** at the intragenic 84 bp interval *ade6-M216*×*ade6-3083*: UoA110×UoA100 (WT, n = 12), UoA478×UoA476 (*msh2-30*, n = 6), UoA494×UoA492 (*msh3*Δ, n = 6), UoA482×UoA480 (*msh6*Δ, n = 6), UoA364×UoA361 (*mlh1*Δ, n = 8), UoA407×UoA405 (*pms1-16*, n = 5), UoA828×UoA830 (*msh2-30 mlh1*Δ, n = 6); **(B)** at the intragenic 33 bp interval *ade6-149*×*ade6-3049*: UoA122×UoA497 (WT, n = 6), UoA368×UoA512 (*mlh1*Δ, n = 6); **(C)** at the intragenic 53 bp interval *ade6-3049*×*ade6-51*: UoA120×UoA463 (WT, n = 6), UoA366×UoA511 (*mlh1*Δ, n = 6); **(D)** at the intragenic 1,320 bp interval *ade6-3083*×*ade6-469*: ALP733×ALP731 (WT, n = 20), UoA477×UoA479 (*msh2-30*, n = 6), UoA493×UoA495 (*msh3*Δ, n = 6), UoA481×UoA483 (*msh6*Δ, n = 6), UoA362×UoA371 (*mlh1*Δ, n = 11), UoA406×UoA410 (*pms1-16*, n = 6), UoA827×UoA829 (*msh2-30 mlh1*Δ, n = 6); **(E)** at the intragenic 1,335 bp interval *ade6-M375*×*ade6-469*: ALP1541×ALP731 (WT, n = 16), UoA510×UoA371 (*mlh1*Δ, n = 6); **(F)** at the intragenic 1,168 bp interval *ade6-M216*×*ade6-3049*: UoA99×UoA123 (WT, n = 12), UoA368×UoA361 (*mlh1*Δ, n = 12); n indicates the number of independent crosses. For details of data see Supplementary Table S3.

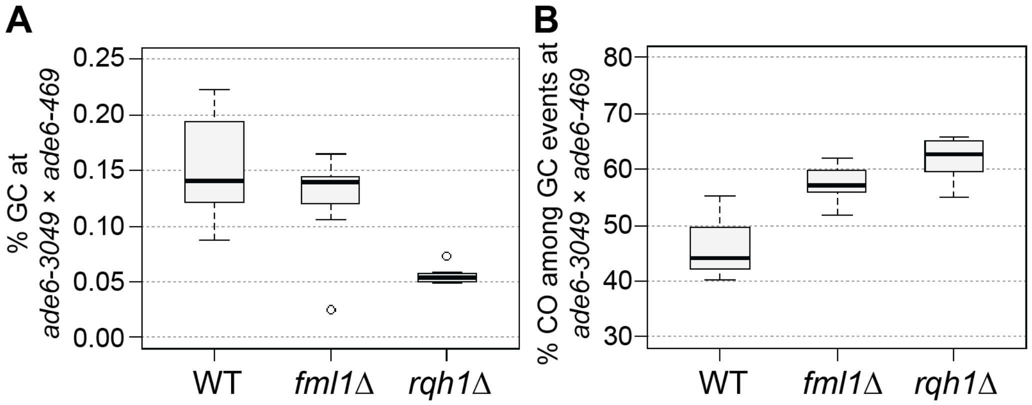


**Supplementary Figure S5.** Rqh1 and Fml1 modulating meiotic recombination outcome at the intragenic 254 bp interval *ade6-3049*×*ade6-469*: **(A)** Frequency of gene conversion (GC) in wild type (WT), *fml1*, and *rqh1* mutants, UoA120×ALP731 (WT, n = 31), ALP1716×MCW4718 (*fml1*Δ, n = 11), MCW6587×ALP780 (*rqh1*Δ, n = 10); **(B)** Frequency of crossovers (CO) among GC events at *ade6* in wild type (WT), *fml1*, and *rqh1* mutants, crosses as in (A). n indicates the number of independent crosses. For details of data see Supplementary Table S3.

**Supplementary Table S1.** Sequence and position (counted from the A of the start codon ATG as first position) of *ade6* point mutations (indicated in bold)

| allele | position | DNA sequence | reference |
|---|---|---|---|
| *ade6-M216* | G47A | ggtcaattgg**A**ccgaatgatg | Szankasi *et al.*, 1988[3] |
| *ade6-M375* | G133T | acaaattgat**T**gaggacgtga | Szankasi *et al.*, 1988[3] |
| *ade6-M26* | G136T | aattgatgga**T**gacgtgagca | Szankasi *et al.*, 1988[3] |
| *ade6-3074* | G136T/G142C | aattgatgga**T**gacgt**C**agcacattga | Steiner & Smith, 2005[4] |
| *ade6-3083* | A131G/G134T/G136T/G142C /G144T/A146G/A148C | aaattg**G**tg**T**a**T**gacgt**C**a**T**c**G**c**C**ttgatgc | Steiner & Smith, 2005[4] |
| *ade6-704*[a] | T645A | ataatgtttg**A**catttagtat | Park *et al.*, 2007[5] |
| *ade6-52*[b] | G796A | tttactcaac**A**aaattgctcc | Steiner *et al.*, 2009[6] |
| *ade6-149* | C1181T | atcatgggtt**T**ggattctgat | Schär & Kohli, 1993[7] |
| *ade6-3049* | C1214A | aaagatgctg**A**cgtcatttta | Steiner & Smith, 2005[4] |
| *ade6-51* | C1267T | tgtttcagct**T**accgcacacc | Schär *et al.*, 1993[8] |
| *ade6-469* | C1468T | tcagatgcct**T**gaggtgtccc | Szankasi *et al.*, 1988[3] |

[a]previously estimated by positional mapping to be C846A[7]; theoretically both, T645A and C846A, create a UGA stop codon suppressible by *sup3-5*[5].
[b]previously reported as T956C[8]

**Supplementary Table S4.** Yeast strain list

| Strain | Relevant genotype | Origin |
|---|---|---|
| ALP729 | *h$^{+S}$ arg3-D4 his3-D1 leu1-32 ura4-D18* | lab strain[9] |
| ALP731 | *h$^{-smt0}$ ade6-469 his3$^{+}$-aim arg3-D4 his3-D1 ura4-D18* | Lorenz *et al.*, 2010[10] |
| ALP733 | *h$^{+S}$ ade6-3083 ura4$^{+}$-aim2 his3-D1 leu1-32 ura4-D18* | Lorenz *et al.*, 2010[10] |
| ALP780 | *h$^{-smt0}$ rqh1Δ::kanMX6 ade6-469 his3$^{+}$-aim arg3-D4 his3-D1 ura4-D18* | Lorenz *et al.*, 2014[11] |
| ALP781 | *h$^{+S}$ rqh1Δ::kanMX6 ade6-3083 ura4$^{+}$-aim2 his3-D1 leu1-32 ura4-D18* | Lorenz *et al.*, 2014[11] |
| ALP1133 | *h$^{+S}$ fml1Δ::hphMX4 ade6-3083 ura4$^{+}$-aim2 his3-D1 leu1-32 ura4-D18* | Lorenz *et al.*, 2012[9] |
| ALP1541 | *h$^{+N}$ ade6-M375 ura4$^{+}$-aim2 his3-D1 leu1-32 ura4-D18* | Lorenz *et al.*, 2012[9] |
| ALP1594 | *h$^{-smt0}$ ade7-50 arg3-D4 his3-D1 ura4-D18* | lab strain[12] |
| ALP1716 | *h$^{+S}$ fml1Δ::hphMX4 ura4$^{+}$-aim2 ade6-3049 his3-D1 leu1-32 ura4-D18* | this study |
| FO652 | *h$^{-smt0}$ arg3-D4 his3-D1 leu1-32 ura4-D18* | lab strain[11] |
| FO1285 | *h$^{+N}$ ade6-M26 ura4$^{+}$-aim2 arg3-D4 his3-D1 leu1-32 ura4-D18* | lab strain |
| MCW4718 | *h$^{-smt0}$ fml1Δ::hphMX4 ade6-469 his3$^{+}$-aim arg3-D4 his3-D1 ura4-D18* | Lorenz *et al.*, 2012[9] |
| MCW6587 | *h$^{+S}$ rqh1Δ::kanMX6 ura4$^{+}$-aim2 ade6-3049 his3-D1 leu1-32 ura4-D18* | this study |
| UoA95 | *h$^{+S}$ ura4$^{+}$-aim2 his3-D1 leu1-32 ura4-D18* | this study |
| UoA96 | *h$^{-smt0}$ ura4$^{+}$-aim2 his3-D1 leu1-32 ura4-D18* | this study |
| UoA97 | *h$^{+S}$ his3+-aim arg3-D4 his3-D1 ura4-D18* | this study |
| UoA98 | *h$^{-smt0}$ his3+-aim arg3-D4 his3-D1 ura4-D18* | this study |
| UoA99 | *h$^{+S}$ ade6-M216 ura4$^{+}$-aim2 his3-D1 leu1-32 ura4-D18* | this study |
| UoA100 | *h$^{-smt0}$ ade6-M216 ura4$^{+}$-aim2 his3-D1 leu1-32 ura4-D18* | this study |
| UoA104 | *h$^{+S}$ ade6-3074 ura4$^{+}$-aim2 arg3-D4 his3-D1 ura4-D18* | this study |
| UoA106 | *h$^{+S}$ ade6-3074 his3$^{+}$-aim arg3-D4 his3-D1 ura4-D18* | this study |
| UoA110 | *h$^{+S}$ ade6-3083 his3$^{+}$-aim arg3-D4 his3-D1 ura4-D18* | this study |
| UoA112 | *h$^{+S}$ ade6-704 ura4$^{+}$-aim2 his3-D1 leu1-32 ura4-D18* | this study |
| UoA115 | *h$^{-smt0}$ ade6-704 his3$^{+}$-aim arg3-D4 his3-D1 ura4-D18* | this study |
| UoA116 | *h$^{+S}$ ade6-52 ura4$^{+}$-aim2 his3-D1 leu1-32 ura4-D18* | this study |
| UoA119 | *h$^{-smt0}$ ade6-52 his3$^{+}$-aim arg3-D4 his3-D1 ura4-D18* | this study |
| UoA120 | *h$^{+S}$ ade6-3049 ura4$^{+}$-aim2 his3-D1 leu1-32 ura4-D18* | this study |
| UoA122 | *h$^{+S}$ ade6-3049 his3$^{+}$-aim arg3-D4 his3-D1ura4-D18* | this study |
| UoA123 | *h$^{-smt0}$ ade6-3049 his3$^{+}$-aim arg3-D4 his3-D1 ura4-D18* | this study |
| UoA361[a] | *h$^{-smt0}$ mlh1Δ::kanMX6 ade6-M216 ura4$^{+}$-aim2 his3-D1 leu1-32 ura4-D18* | this study |
| UoA362[a] | *h$^{+S}$ mlh1Δ::kanMX6 ade6-3083 ura4$^{+}$-aim2 his3-D1 leu1-32 ura4-D18* | this study |
| UoA364[a] | *h$^{+S}$ mlh1Δ::kanMX6 ade6-3083 his3$^{+}$-aim arg3-D4 his3-D1 ura4-D18* | this study |
| UoA366[a] | *h$^{+S}$ mlh1Δ::kanMX6 ade6-3049 ura4$^{+}$-aim2 his3-D1 leu1-32 ura4-D18* | this study |
| UoA368[a] | *h$^{+S}$ mlh1Δ::kanMX6 ade6-3049 his3$^{+}$-aim arg3-D4 his3-D1 ura4-D18* | this study |
| UoA371[a] | *h$^{-}$ mlh1Δ::kanMX6 his3$^{+}$-aim ade6-469 arg3-D4 his3-D1 ura4-D18* | this study |
| UoA405[b] | *h$^{-smt0}$ pms1-16::natMX4 ade6-M216 ura4$^{+}$-aim2 his3-D1 leu1-32 ura4-D18* | this study |
| UoA406[b] | *h$^{+S}$ pms1-16::natMX4 ura4$^{+}$-aim2 ade6-3083 his3-D1 leu1-32 ura4-D18* | this study |
| UoA407[b] | *h$^{+S}$ pms1-16::natMX4 ade6-3083 his3$^{+}$-aim arg3-D4 his3-D1 ura4-D18* | this study |
| UoA410[b] | *h$^{-smt0}$ pms1-16::natMX4 his3$^{+}$-aim ade6-469 arg3-D4 his3-D1 ura4-D18* | this study |
| UoA447 | *h$^{-smt0}$ fml1Δ::natMX6 ade6-M216 ura4$^{+}$-aim2 his3-D1 leu1-32 ura4-D18* | this study |
| UoA450 | *h$^{+S}$ fml1Δ::natMX6 ade6-3083 his3$^{+}$-aim arg3-D4 his3-D1 ura4-D18* | this study |
| UoA459 | *h$^{-smt0}$ msh2-30::hphMX4 arg3-D4 his3-D1 leu1-32 ura4-D18* | this study |
| UoA460 | *h$^{+S}$ msh3Δ-32::kanMX6 arg3-D4 his3-D1 leu1-32 ura4-D18* | this study |
| UoA461 | *h$^{-smt0}$ msh6Δ-34::natMX4 arg3-D4 his3-D1 leu1-32 ura4-D18* | this study |

| | | |
|---|---|---|
| UoA463 | *h$^{-smt0}$ ade6-51 his3$^{+}$-aim arg3-D4 his3-D1 ura4-D18* | this study |
| UoA476 | *h$^{-smt0}$ msh2-30::hphMX4 ade6-M216 ura4$^{+}$-aim2 his3-D1 leu1-32 ura4-D18* | this study |
| UoA477 | *h$^{+S}$ msh2-30::hphMX4 ade6-3083 ura4$^{+}$-aim2 his3-D1 leu1-32 ura4-D18* | this study |
| UoA478 | *h$^{+S}$ msh2-30::hphMX4 ade6-3083 his3$^{+}$-aim arg3-D4 his3-D1 ura4-D18* | this study |
| UoA479 | *h$^{-smt0}$ msh2-30::hphMX4 ade6-469 his3$^{+}$-aim arg3-D4 his3-D1 ura4-D18* | this study |
| UoA480 | *h$^{-smt0}$ msh6Δ-34::natMX4 ade6-M216 ura4$^{+}$-aim2 his3-D1 leu1-32 ura4-D18* | this study |
| UoA481 | *h$^{+S}$ msh6Δ-34::natMX4 ade6-3083 ura4$^{+}$-aim2 his3-D1 leu1-32 ura4-D18* | this study |
| UoA482 | *h$^{+S}$ msh6Δ-34::natMX4 ade6-3083 his3$^{+}$-aim arg3-D4 his3-D1 ura4-D18* | this study |
| UoA483 | *h$^{-smt0}$ msh6Δ-34::natMX4 ade6-469 his3$^{+}$-aim arg3-D4 his3-D1 ura4-D18* | this study |
| UoA492 | *h$^{-smt0}$ msh3Δ-32::kanMX6 ade6-M216 ura4$^{+}$-aim2 his3-D1 leu1-32 ura4-D18* | this study |
| UoA493 | *h$^{+S}$ msh3Δ-32::kanMX6 ade6-3083 ura4$^{+}$-aim2 his3-D1 leu1-32 ura4-D18* | this study |
| UoA494 | *h$^{+S}$ msh3Δ-32::kanMX6 ade6-3083 his3$^{+}$-aim arg3-D4 his3-D1 ura4-D18* | this study |
| UoA495 | *h$^{-smt0}$ msh3Δ-32::kanMX6 ade6-469 his3$^{+}$-aim arg3-D4 his3-D1 ura4-D18* | this study |
| UoA497 | *h$^{-smt0}$ ade6-149 ura4$^{+}$-aim2 his3-D1 leu1-32 ura4-D18* | this study |
| UoA499 | *h$^{-smt0}$ rqh1Δ-G1::natMX6 ade6-M216 ura4$^{+}$-aim2 his3-D1 leu1-32 ura4-D18* | this study |
| UoA502 | *h$^{+S}$ rqh1Δ-G1::natMX6 ade6-3083 his3$^{+}$-aim arg3-D4 his3-D1 ura4-D18* | this study |
| UoA510 | *h$^{+N}$ mlh1Δ::kanMX6 ura4$^{+}$-aim2 ade6-M375 his3-D1 leu1-32 ura4-D18* | this study |
| UoA511[a] | *h$^{-smt0}$ mlh1Δ::kanMX6 ade6-51 his3$^{+}$-aim arg3-D4 his3-D1 ura4-D18* | this study |
| UoA512[a] | *h$^{-smt0}$ mlh1Δ::kanMX6 ade6-149 ura4$^{+}$-aim2 his3-D1 leu1-32 ura4-D18* | this study |
| UoA827[a] | *h$^{+S}$ mlh1Δ::kanMX6 msh2-30::hphMX4 ura4$^{+}$-aim2 ade6-3083 his3-D1 leu1-32 ura4-D18* | this study |
| UoA828[a] | *h$^{+S}$ mlh1Δ::kanMX6 msh2-30::hphMX4 ade6-3083 his3$^{+}$-aim arg3-D4 his3-D1 ura4-D18* | this study |
| UoA829[a] | *h$^{+S}$ mlh1Δ::kanMX6 msh2-30::hphMX4 his3$^{+}$-aim ade6-3083 arg3-D4 his3-D1 ura4-D18* | this study |
| UoA830[a] | *h$^{-smt0}$ mlh1Δ::kanMX6 msh2-30::hphMX4 ade6-M216 ura4$^{+}$-aim2 his3-D1 leu1-32 ura4-D18* | this study |
| UoA861 | *h$^{+S}$ ade6-M375 his3$^{+}$-aim arg3-D4 his3-D1 ura4-D18* | this study |

[a]*mlh1Δ* strains are derivatives of OL937[13], provided as FY18813 by the National BioResource Project (NBRP) of the MEXT, Japan.

[b]*pms1* insertion mutant strains are derivatives of PRS301[14], provided as FY18790 by the National BioResource Project (NBRP) of the MEXT, Japan.
